# Ribosome-profiling reveals restricted post transcriptional expression of antiviral cytokines and transcription factors during SARS-CoV-2 infection

**DOI:** 10.1101/2021.03.03.433675

**Authors:** Marina R. Alexander, Aaron M. Brice, Petrus Jansen van Vuren, Christina L. Rootes, Leon Tribolet, Christopher Cowled, Andrew G. D. Bean, Cameron R. Stewart

## Abstract

The global COVID-19 pandemic caused by SARS-CoV-2 has resulted in over 2.2 million deaths. Disease outcomes range from asymptomatic to severe with, so far, minimal genotypic change to the virus so understanding the host response is paramount. Transcriptomics has become incredibly important in understanding host-pathogen interactions; however, post-transcriptional regulation plays an important role in infection and immunity through translation and mRNA stability, allowing tight control over potent host responses by both the host and the invading virus. Here we apply ribosome profiling to assess post-transcriptional regulation of host genes during SARS-CoV-2 infection of a human lung epithelial cell line (Calu-3). We have identified numerous transcription factors (JUN, ZBTB20, ATF3, HIVEP2 and EGR1) as well as select antiviral cytokine genes, namely IFNB1, IFNL1,2 and 3, IL-6 and CCL5, that are restricted at the post-transcriptional level by SARS-CoV-2 infection and discuss the impact this would have on the host response to infection. This early phase restriction of antiviral transcripts in the lungs may allow high viral load and consequent immune dysregulation typically seen in SARS-CoV-2 infection.

## INTRODUCTION

Coronaviruses are enveloped positive sense RNA viruses with an exceptionally large genome encoding the structural proteins envelope (E), spike (S), membrane (M), and nucleocapsid (N), in addition to a plethora of non-structural and accessory proteins. Infections with endemic human coronaviruses (e.g. 229E, NL63, OC43, and HKU1) cause a mild common cold, however three novel coronaviruses have emerged from animal reservoirs in the 21^st^ century, SARS-CoV, MERS and SARS-CoV-2 causing a fatal respiratory syndrome in 34%, 15%, and 3% of cases, respectively with SARS-CoV-2 being the most infectious [1]. The current COVID-19 pandemic caused by SARS-CoV-2 emerged in China in December 2019 [2] and has since spread across the globe, causing more than 100 million confirmed cases and over 2 million deaths (https://covid19.who.int/ accessed February 2021).

SARS-CoV-2 is adept at evading innate immunity [3], the naïve host’s primary defense against a newly emerged coronavirus. It does this using numerous structural and non-structural proteins that inhibit interferon (IFN) production and function. Through blocking IFN, viral replication can proceed unchecked. In addition to antiviral functions, circulating IFN alerts the host to viral infection with symptoms such as fever, pain and fatigue [4]. This capacity for stealth replication in the absence of symptoms allows asymptomatic transmission [5], making it particularly difficult to control from a public health perspective. Understanding the molecular mechanics of immune evasion strategies employed by SARS-CoV-2 will allow us to develop more effective diagnostics, therapeutics, and host prognostic indicators.

RNA sequencing (RNA-seq) enables discovery of molecular pathways involved in SARS-CoV-2 infection. RNA-seq of nasopharyngeal swabs from infected individuals revealed up-regulation of antiviral factors OAS1-3 and IFIT1-3 as well as chemokines like IP-10, the latter of which showed muted expression in older individuals [6]. Added to this, transcriptional profiles in SARS-CoV-2 infected cell, animal and patient serum samples revealed low levels of type I and III interferons but high levels of TNF, IL-6, RANTES and CCL20 compared to influenza A infection, indicating differences in the immune responses to SARS-CoV-2 compared to another respiratory virus [7]. These studies have provided a greater insight into potential mechanisms underpinning COVID-19 immunopathogenesis. However, they do not capture post-transcriptional regulatory mechanisms, many of which control the initiation, magnitude, duration, and resolution of the innate immune response [8] or allow the virus to restrict host gene expression [9].

Here we have used ribosome profiling to assess post-transcriptional regulation of host gene expression during SARS-CoV-2 infection. We found restricted translation of type I and III IFNs 24 hours post infection as well as numerous transcription factors and antiviral cytokines. This data provides a set of genes likely responsible for the delayed antiviral responses seen in SARS-type coronavirus infections strengthening development of molecular-based strategies to combat this devastating pandemic.

## RESULTS

### Development of a SARS-CoV-2 infection model in Calu-3 cells

Of the multiple organs that SARS-CoV-2 infects, viral load is highest in the lungs, and infection of this organ is a significant driver of pathogenesis [10]. We therefore chose a lung epithelial cell line known to be productively infected with SARS-CoV-2 to investigate host responses to infection. Calu-3 cells are a human immortalized cell line derived from lung adenocarcinoma which express sufficient levels of ACE2 to allow SARS-Cov-2 entry. We found SARS-CoV-2 infectious titres peaked 24 hours post infection in Calu-3 cells (Figure 1A) as per other studies [11]. Both SARS-CoV-2 intracellular genomic RNA (Figure 1B) and N protein (Figure 1C) followed similar kinetics with peak viral load post 24 hours infection. Immunofluorescence staining of SARS-CoV-2 N-protein over a time-course from 1-48hrs showed infection of more than 50% of cells after 24 hours (Figure 1D).

**Figure 1.**
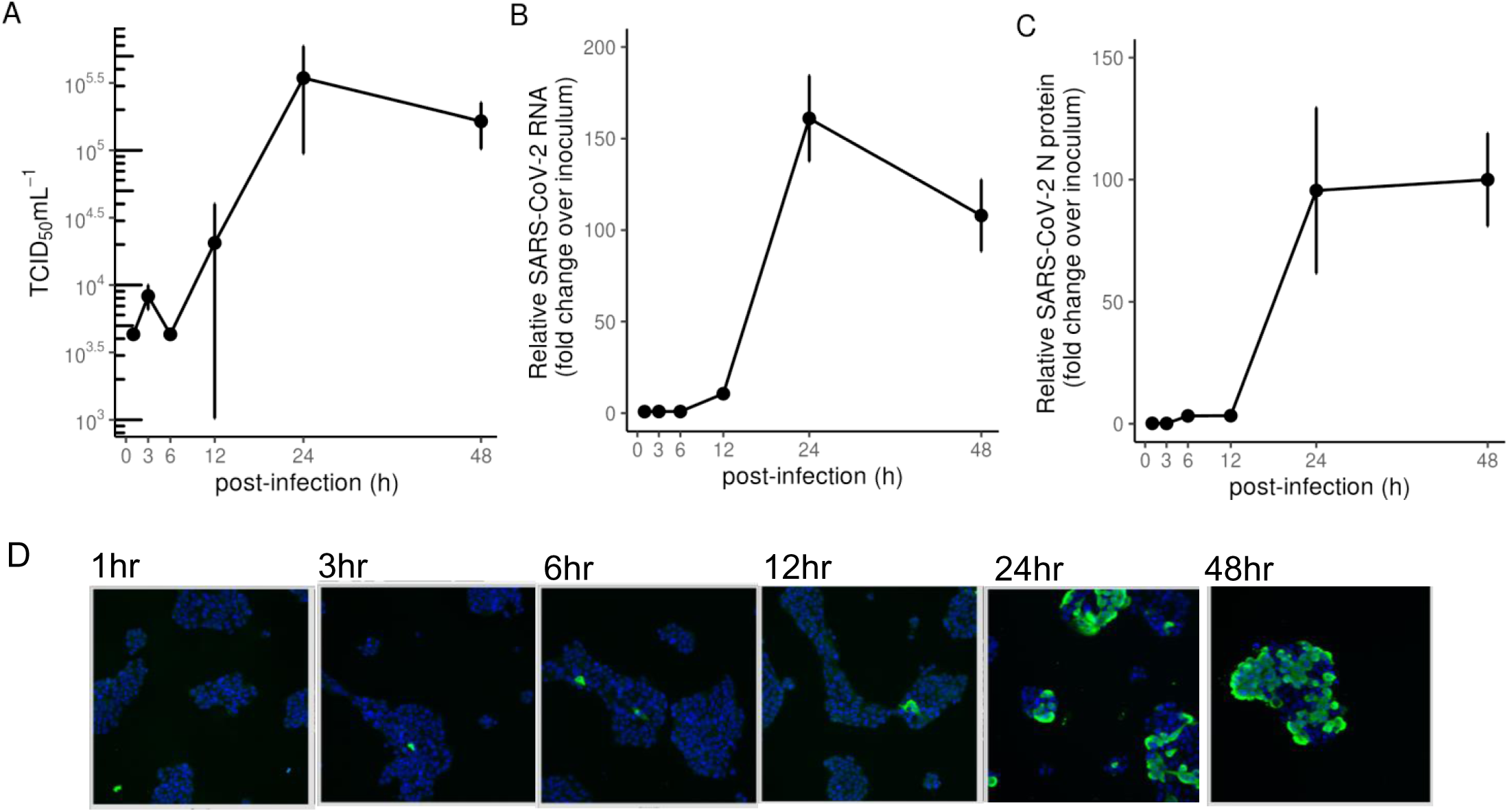
SARS-CoV-2 infection of Calu-3 cells. (A) TCID_50_ measurements of virus titres, (B) qRT-PCR measurements of intracellular viral RNA represented by 2^-ΔΔCt^ normalised first to *GAPDH* and then to inoculum levels of SARS-CoV-2, set to 1, and (C) intracellular viral protein in Calu-3 cells infected with SARS-CoV-2 (MOI 0.1). (D) Immunofluorescence microscopy showing SARS-CoV-2 N protein staining (green) in Calu-3 cells infected with SARS-CoV-2 (MOI 0.1) at various timepoints. Cell nuclei were stained using DAPI (blue).

Having developed an infection model in Calu-3 cells, we proceeded to capture transcriptional and post-transcriptional changes in gene expression at 24 hours post SARS-CoV-2 infection using a method deemed suitable for high containment infections [12]. Cells were snap frozen in liquid nitrogen to lock ribosomes onto their associated mRNA, avoiding artifacts introduced by translation inhibitors like cycloheximide [13]. Both total RNA and RNA protected by protein from micrococcal nuclease (MNase) digest were purified. The latter, named herein as ribosome footprints were further purified to contain fragments between 25 and 35nt which spans the possible 80S ribosomal footprint on actively translated mRNAs (Figure A1). Approximately 30 million reads were obtained from next generation sequencing of the total mRNA libraries, while more than 50 million reads were obtained from the MNase-digested small RNA libraries.

### Transcriptional response to SARS-CoV-2 infection of Calu-3 cells is dominated by antiviral defence genes

We first analysed reads obtained from total RNA representing the transcriptome. 11504 genes passed the threshold of >1 counts per million in at least three samples and were analysed for differential expression by comparing triplicate mock and SARS-CoV-2-infected samples. There were 2.5 times more up-regulated than down-regulated genes (166 versus 63, Figure 2A) using a log2 fold change cut-off of 1 and p-value cut-off of 0.05. Upon inspection of the genes up-regulated by SARS-CoV-2 infection, many genes were previously annotated antiviral genes (Figure 2A). To confirm this, the 229 significantly altered genes were submitted to https://david.ncifcrf.gov/ for functional annotation clustering using the Uniprot keyword database. The top seven significantly enriched keywords are plotted in Figure 2B. Antiviral defence showed the highest fold enrichment and a p-value <0.001. Other significantly enriched keywords included innate immunity, RNA-binding and transcription. Of the 134 Antiviral defence keywords, 27 were up-regulated by SARS-CoV-2 and are listed in Figure 2C. Many of these genes, including IFIT1, IFIT2, IFIT3, RSAD2, OASL, HERC5, DDX58 and MX2 were also upregulated in nasopharyngeal swabs of COVID19 patients [6].

**Figure 2.**
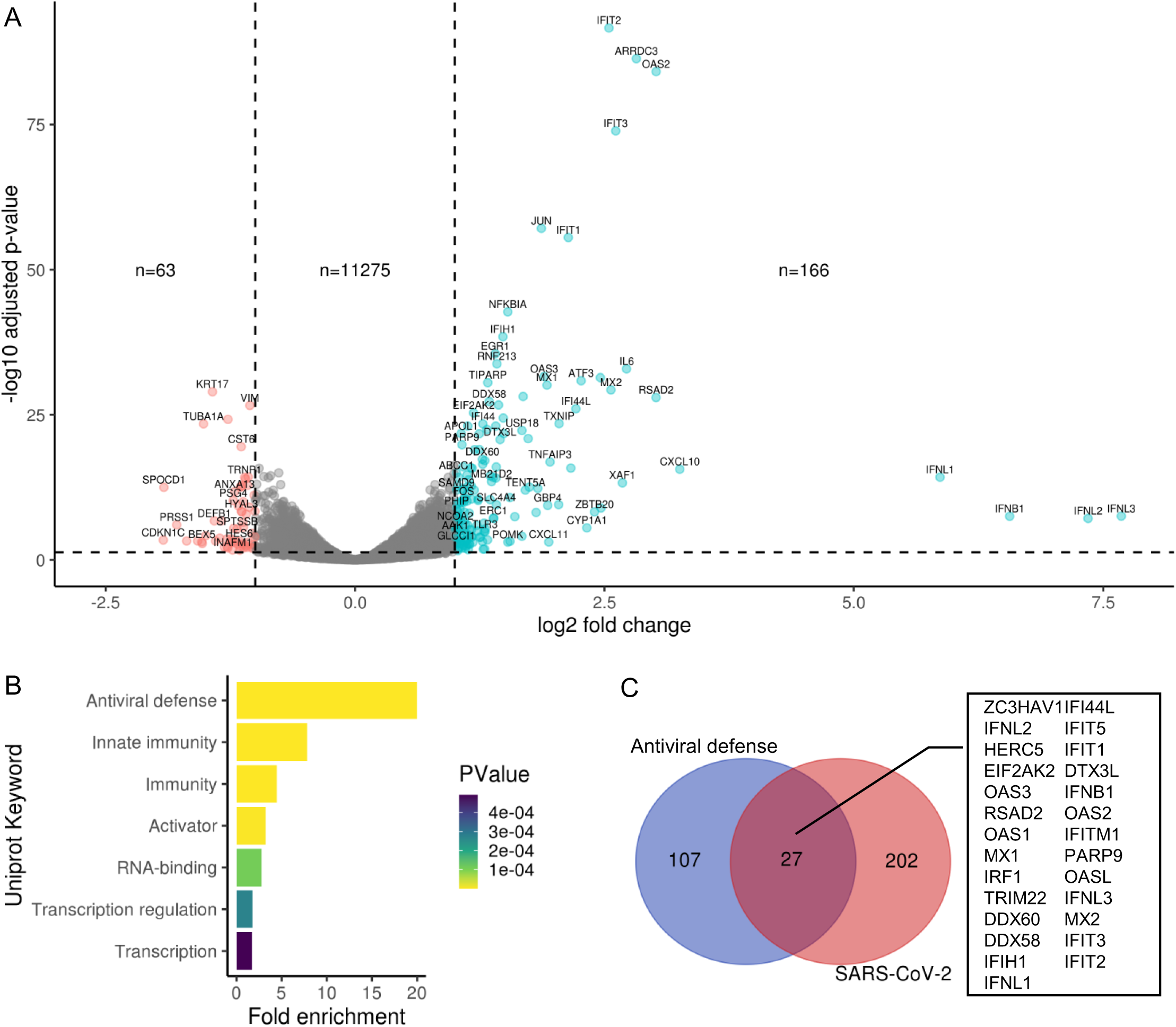
Transcriptional response to SARS-CoV-2 infection of Calu3 cells is dominated by antiviral defense genes. (A) Volcano plot showing global transcriptional changes of ∼11,000 genes in SARS-CoV-2 infected Calu-3 cells. Log2FoldChange infected 24 hours versus mock. 229 transcripts were differentially expressed (DE) in SARS-CoV-2 infected cells based on the cut-off of p-value <0.05, log2fold change >1. (B) Significantly altered genes were submitted to https://david.ncifcrf.gov/ for Functional Annotation Clustering. Here Uniprot Keywords were summarised using fold enrichment of DE genes, coloured by PValue. (C) Common genes between Uniprot “Antiviral Defense” keywords and the 229 SARS-CoV-2 DE genes.

### MNAse digestion of cell extracts yields CDS-mapped ribosome footprints

Next, we proceeded to analyse the reads generated from sequencing of size-selected MNase-digested RNA to understand the translatome in SARS-CoV-2 infected cells. 50% of the reads (approximately 25 million reads) mapped to protein coding sequence in both mock and infected cells (Figure 3A), indicating close to 50% coverage of the 18 million nucleotides of coding sequence predicted in the human transcriptome [14]. This finding implies that sequencing depth was sufficient to make gene level inferences on differential ribosome density. More than 50% of reads mapped to the first 30 nucleotides of tRNAs mostly carrying the small neutral amino acid Glycine and some carrying the small to medium sized acidic amino acids, Glutamate and Aspartate (data not shown). These tRNA reads and the less abundant reads mapping to other non-coding RNA were filtered out and remaining reads mapped to different features of human protein coding genes. Most of these filtered reads (60%) mapped to within the CDS (Figure 3B). The 3’UTR and 5’UTR had 20% and 7% of the reads respectively no significant difference between mock and infected cells was observed. Annotated start and stop codons were present in 2% and 6% of the reads, respectively with no difference between mock and infected. When high-resolution metagene analysis of transcript coverage was performed, a gradual increase in coverage was seen leading up the start codon, whereby coverage increased 8-fold to a peak around 30nt consistent with ribosomes initiating at the start codon (Figure 3C and D).

**Figure 3.**
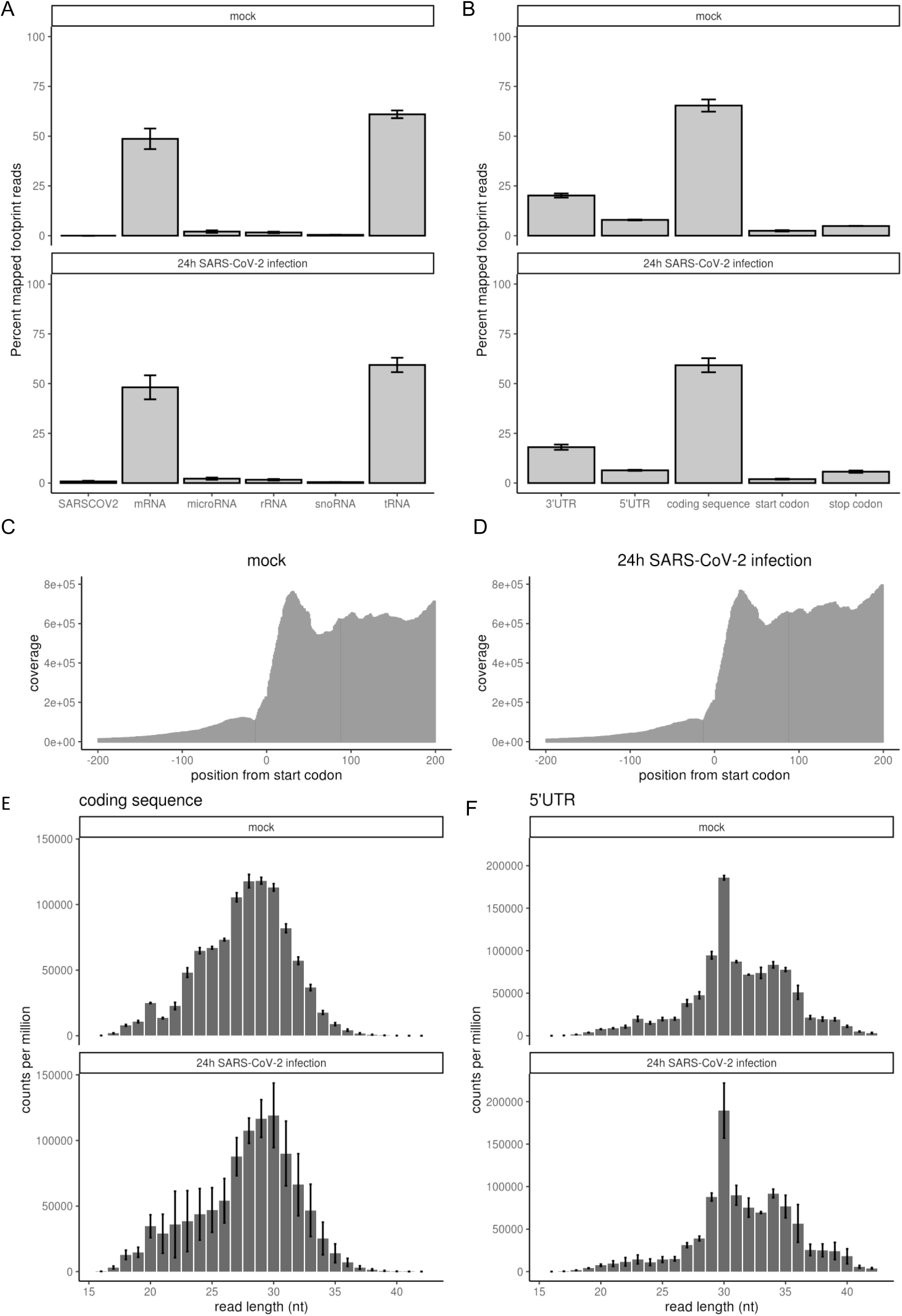
MNAse digestion of cell extracts yields 30nt long CDS-mapped ribosome footprints. (A) Percentage of reads mapping to indicated RNA species in mock and SARS-CoV-2 infected cells at 24 hours. (B) Percentage of non-coding RNA filtered reads mapping to indicated gene-level features in mock and SARS-CoV-2 infected cells. (C-D) Metagene coverage aligned to the start codon of all annotated protein coding genes for mock and SARS-CoV-2 infected cells. (E-F) Read-length counts per million read distribution in coding sequence and 5’ untranslated regions (UTR).

Eukaryotic 80S ribosome footprints are ∼28-30 nucleotides in length [15]. To confirm that CDS-mapped reads were bona fide ribosome footprints, read-lengths were calculated for reads mapping to the CDS and the 5’UTR region as 80S ribosomes scan this region to find the start codon [16]. Read-length distributions were summarised across all genes. Density peaked at 28-30 nucleotides in coding sequence in both mock and infected cells indicative of bona-fide ribosome footprints (Figure 3E). Read-lengths higher than 30 nucleotides showed an exponential shortening towards 40 nucleotides suggesting single nucleotide degradation, while those shorter than 30 nucleotides were staggered suggestive of footprints derived from smaller non-ribosomal RNA binding proteins. In the 5’ UTR read lengths also peaked at 30 nucleotides indicative of translation in the 5’ leader as documented elsewhere [17] but at >6-fold lower density compared to the CDS mapped reads (Figure 3E). There was no significant difference in 5’ UTR ribosome occupancy between mock and infected cells apart from greater variance between replicates in the infected samples.

### Transcription factors and cytokines are blocked at the post-transcriptional level

Using CDS-mapped read counts from MNase-digested RNA libraries, we compared the translatome of mock versus SARS-CoV-2 infected Calu3 cells. Here 11,455 genes passed the threshold (>1 counts per million in at least three samples), comparing well to numbers passing threshold expression for the transcriptome (Figure 4A). Three genes (KRT17, CDKN1A and TOM1) showed more than a 2-fold decrease in expression. 18 genes showed more than a 2-fold increase in expression upon infection and are mostly antiviral, i.e., Interferon-induced proteins with tetratricopeptide repeats (IFIT), oligoadenylate synthase (OAS), DDX58 (RIGI) and Interferon-induced GTP-binding protein Mx1 (MX1). Of note is that many genes seen upregulated at the whole transcript level were not seen to be differentially expressed when ribosome-associated RNA was sequenced.This transcriptome-only change contrasts with other stimulatory contexts where 9% of genes showed transcriptome-only upregulation and 85% of genes showed translation-only upregulation [18].

**Figure 4.**
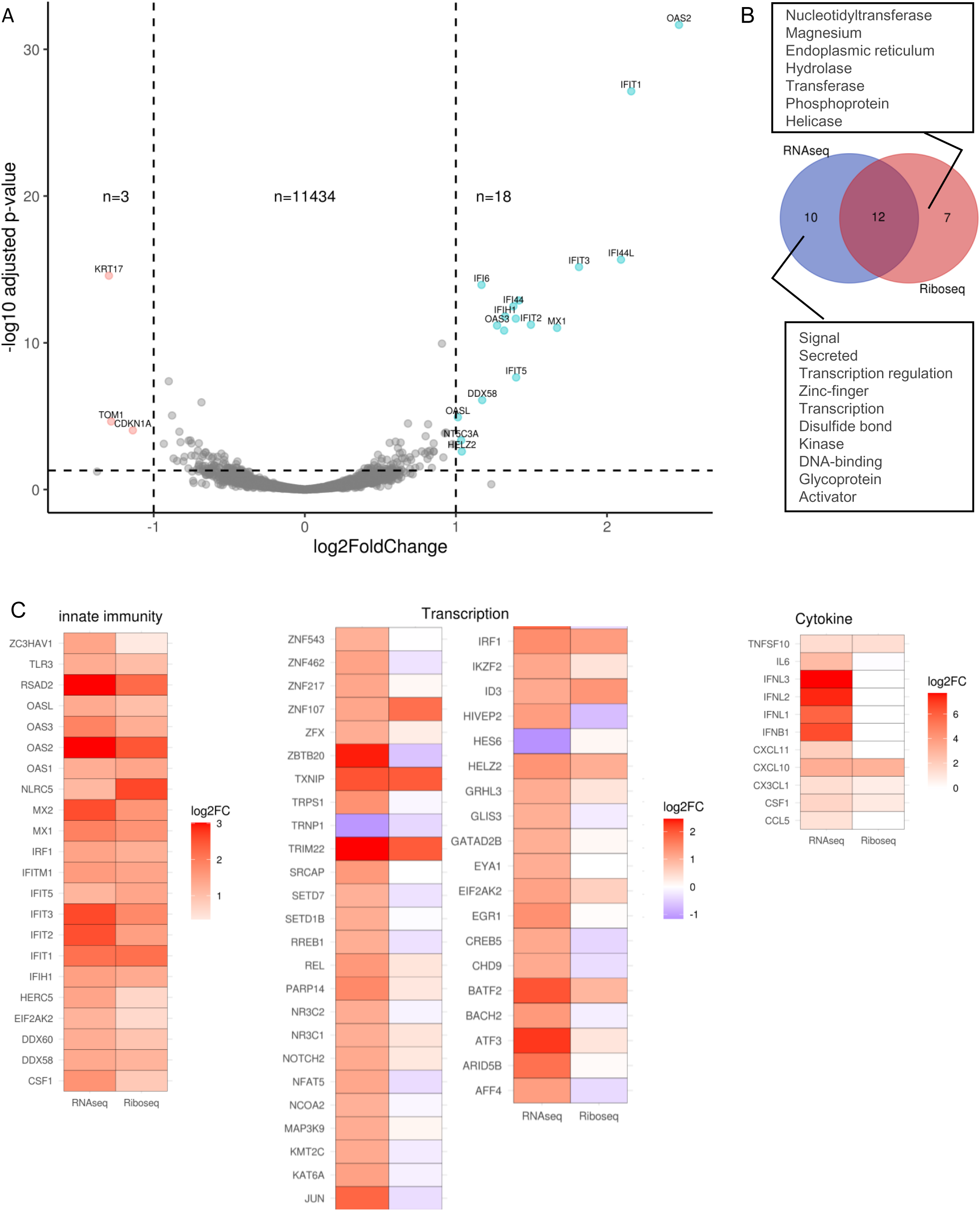
Absence of transcription factors and cytokines from genes up-regulated by ribosome foot-printing. (A) Volcano plot showing global translational changes of ∼11,000 genes in SARS-CoV-2 infected Calu-3 cells. Log2FoldChange infected 24 hours versus mock. 21 genes were differentially expressed (DE) in SARS-CoV-2 infected cells based on the cut-off of p-value <0.05, log2fold change >1. (B) Functional annotation clustering on the differential expression gene lists for RNA-seq versus Ribo-seq. Uniprot keywords used to classify genes showed an absence of keywords ‘Signal’, ‘Secreted’ and ‘Transcription regulation’ in the Ribo-seq gene list. (C) Heat-map of the log2 fold change of up-regulated RNA-seq transcripts included within ‘Innate immunity’, ‘Transcription’ and ‘Cytokine keywords’ alongside their log2 fold change as determined by Ribo-seq.

We investigated this paucity of up-regulated genes further, using functional annotation clustering on the differential expression gene lists for RNA-seq versus Ribo-seq. Using Uniprot keywords to classify genes, showed an absence of keywords ‘Signal’, ‘Secreted’ and ‘Transcription regulation’ in the Ribo-seq gene list. These three annotation groups include genes such as IFNL1, CCL5, IFNB1, IL6, ATF3, JUN, SETD1B and REL (Figure 4B). To summarise this apparent block in expression at the Ribo-seq level we heat-mapped the log2 fold change of up-regulated RNA-seq transcripts included within ‘Innate immunity’, ‘Transcription’ and ‘Cytokine keywords’ alongside their log2 fold change as determined by Ribo-seq. Within the ‘Innate immunity’ annotation group, most genes upregulated by RNA-seq were also upregulated by Ribo-seq, except for NLRC5, a negative regulator of NFKB and IFN-I signalling [19], which was 2.2-fold and 6.2-fold induced by viral infection for RNA-seq and Ribo-seq, respectively. Within the ‘Transcription’ annotation group, 37 (84%) of the differentially expressed genes by RNA-seq were restricted in expression by Ribo-seq, with one of the most restricted being JUN (3.7-fold increase by RNA-seq and 1.3-fold decreased by Ribo-seq). Those un-restricted at the Ribo-seq level included ZNF107, TXNIP, IRF1 and TRIM22. The third annotation group we analysed was ‘Cytokine’, where all three Interferon lambdas (IFNL1, IFNL2, IFNL3), Interferon beta, IL6, CCL5 and CXCL11 were not increased at the Ribo-seq level as they were for RNA-seq. Cytokines with no restriction at the Ribo-seq level were TNFSF10 (TRAIL), CXCL10 (IP10), CX3CL1 and CSF1.

### Coverage of individual genes confirms post-transcriptional restriction of IL6, CCL5, JUN and IFNB1

To examine ribosome occupancy pattern across the length of interesting genes we collected sample normalized, per nucleotide coverage statistics for individual transcripts. Coverage values for all transcript variants were summed at each position from the transcription start site. Gene-level coverage for RNA-seq reads was even across the 5’UTR, CDS and 3’UTR with a decrease observed at the 3’-end termini due to terminal exon-skipped transcript variants (Figure 5). Conversely, Ribo-seq coverage was minimal >100 nucleotides up and down from the start and stop codon, respectively, and this was most evident in genes that were well-occupied by ribosomes, i.e. TNFSF10. IRF7 showed noticeable Ribo-seq coverage in the full length of the 5’ and 3’UTRs which were uniquely short. Genes presented here that were upregulated when expression was assayed by RNA-seq and not by Ribo-seq included IL6, JUN, IFNB1 and CCL5 (RANTES). These genes show a noticeable increase in RNA-seq coverage over the CDS from mock to 24h SARS-CoV-2 infection (top right quadrant, Figure 5) consistent with log2 fold change calculated from count data. The transcription factor REL, showed equivalent up-regulation by RNA-seq and Ribo-seq coverage contrary to the count data, which showed a 2.5-fold increase by RNA-seq (p-value = 2.267×10^-7^) versus 1.3-fold increase by Ribo-seq (not significant). Unlike the aforementioned genes; IRF7, CXCL10, TNFSF10 (TRAIL), OAS2 and IRF1 showed up-regulation by RNA-seq and Ribo-seq. This was not due to higher levels of peak coverage as CXCL10 only reached 30 stacked reads per nucleotide (coverage), equivalent to CCL5 which was not concordant between RNA-seq and Ribo-seq in relative changes between mock and 24h SARS-CoV-2 infected. Neither were Ribo-seq reads in the 3’UTR associated with discordance as both IL6 and IRF1 showed peaks >250 nucleotides downstream of the reference sequence stop codon. In summary, while transcript levels for genes like IL6, CCL5, IFNB1 and JUN were upregulated, these transcripts were not increased in their association with ribosomes suggesting a restriction in the antiviral response.

**Figure 5.**
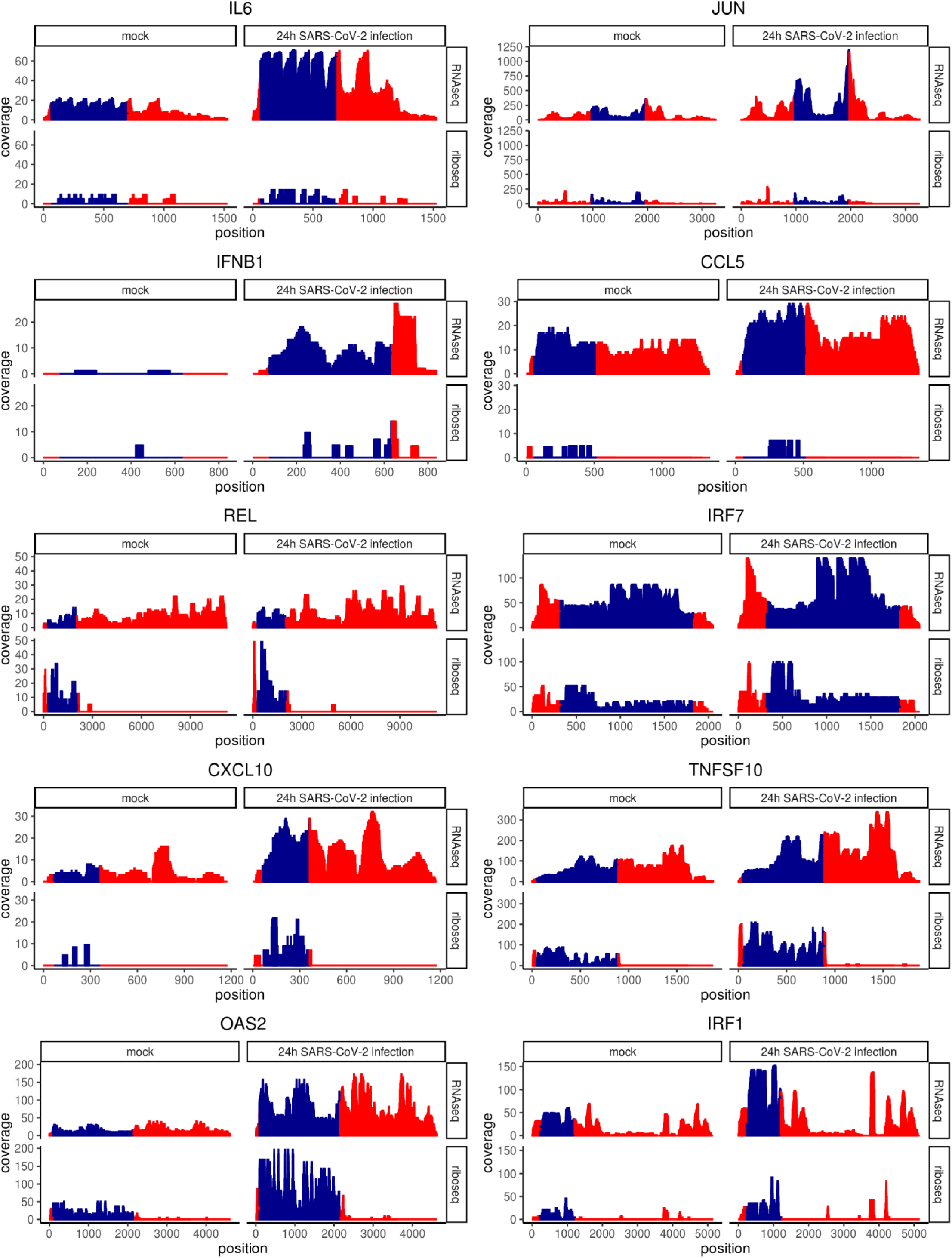
Coverage of select genes confirms post-transcriptional restriction of IL6, JUN, IFNB1 and CCL5. Sample normalized, per nucleotide coverage statistics for indicated transcripts. Coverage values for all transcript variants were summed at each position from the transcription start site. CDS cordinates (blue) were determined using the default transcript per gene as outlined in the Matched Annotation from NCBI and EMBL-EBI (MANE) project. Red indicates 5’ and 3’ untranslated regions.

### mRNA features influence the likelihood of translation inhibition

Differential expression analysis of Ribo-seq count data allowed us to examine the genes which up-regulated in ribosomal occupancy and is a closer approximation to protein output and phenotype than RNA-seq [20]. However, to understand the mechanism through which a restriction in ribosome occupancy occurs, we need to separate the two processes of transcription and translation. Translation efficiency considers the background mRNA levels. We utilized a method, Riborex, whereby RNA-seq count data obtained from matched biological samples is factored into a generalized linear model for the Ribo-seq counts to find genes with differential translational efficiency [21].

Many transcription factors and secreted proteins, which encompass cytokines were found to have short half-lives in mouse embryonic stem cells [22]. To assist in explaining why some cytokines and transcription factors were sensitive to translation inhibition while others were not, we categorized genes based on their mRNA stability as documented by Sharova et al. [22] 6,878 genes with known mRNA half-lives and which passed the default threshold values for Riborex differential expression analysis are presented in Figure 6A. By separating genes into stable mRNAs (half-life >5 hours) and unstable mRNAs (half-life < 5 hours) we found 0.5% of stable mRNAs and 0.2% of unstable mRNAs were increased in translation efficiency. Conversely, more unstable mRNA genes were decreased in translation efficiency, 3.5 % compared to 0.74% of stable mRNAs. The genes with the greatest log2 fold decrease, JUN, ZBTB20 and IL6 were all unstable mRNAs. Across all genes, unstable (half-life <5 hours) mRNAs showed a log2 fold change in translation efficiency significantly lower than stable mRNAs (one-way analysis of variance [ANOVA], *p* < 0.001; Figure 6B.

**Figure 6.**
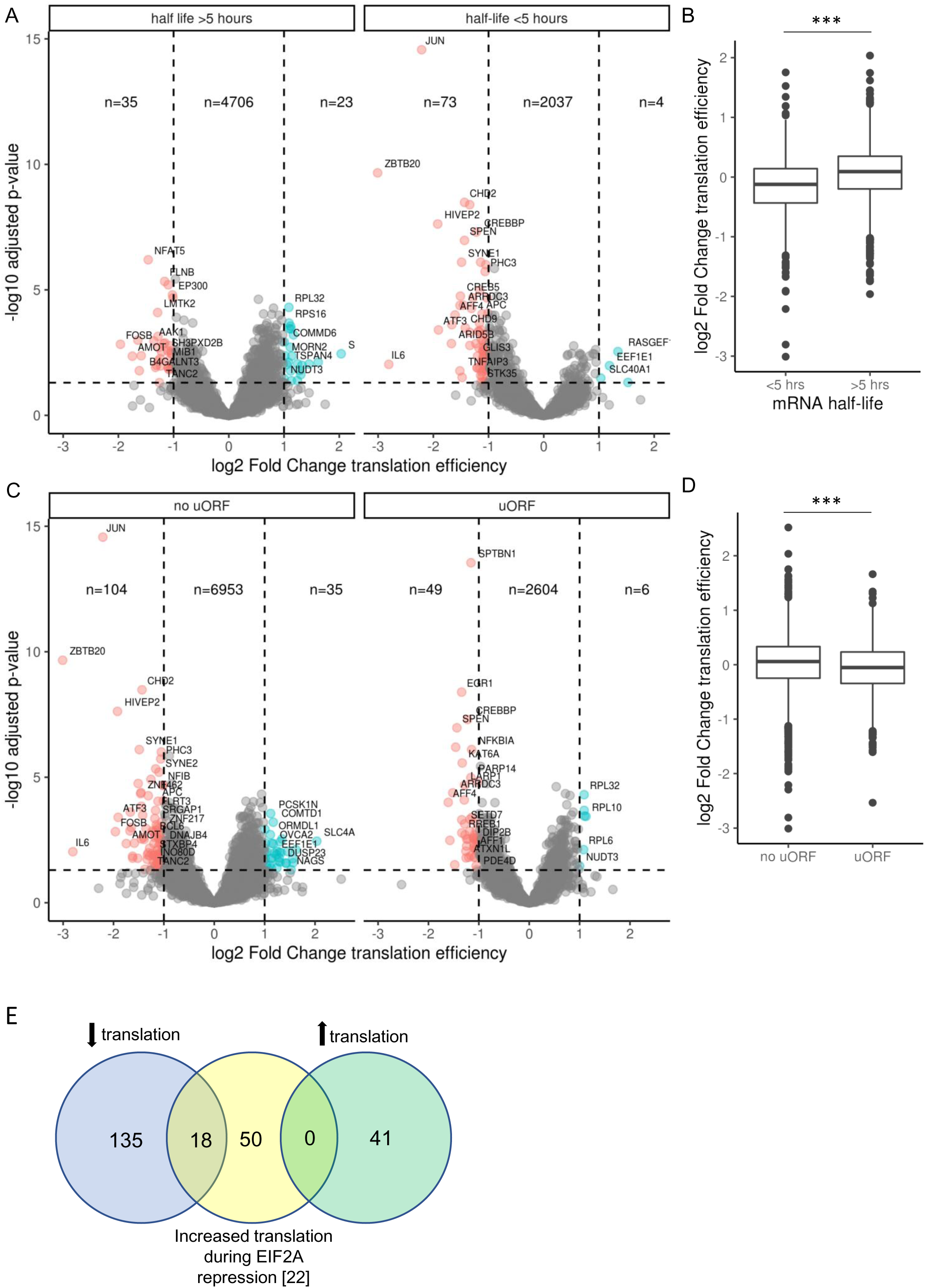
Unstable genes are more sensitive to translation inhibition. **(A)** Volcano plot showing global translation efficiency (Riborex engine) changes of 6,878 genes in SARS-CoV-2 infected Calu-3 cells with previously documented mRNA stability [22]. Log2FoldChange infected 24 hours versus mock. 135 transcripts were differentially expressed (DE) in SARS-CoV-2 infected cells based on the cut-off of p-value <0.05, log2fold change >1. Genes were categorized as having half-life more or less than 5 hours as indicated**. (B)** Log2 fold change as presented in (A) of unstable (<5 hour mRNA half-life) and stable (> 5 hour mRNA half-life) genes presented using geom_boxplot function in the ggplot2 R package using the 25^th^ and 75^th^ percentiles to form the box and whiskers no larger than 1.5 times the interquartile range. Data points beyond the whiskers are outliers. *** indicates one-way analysis of variance [ANOVA], *p* < 0.001. **(C)** Volcano plot showing global translation efficiency changes of 9,751 genes in SARS-CoV-2 infected Calu-3 cells, 2659 of which have one or more high-confidence upstream open reading frames (uORFS) [17]. **(D)** Log2 fold change for genes with no uORF or one or more uORF as presented in (C) using geom_boxplot as described in (B). *** indicates one-way analysis of variance [ANOVA], *p* < 0.001. **(E)** Venn diagram showing genes significantly decreased in translation efficiency by 24 hours of SARS-CoV-2 infection in Calu3 cells (blue), genes increased in translation during EIF2A repression [24] (yellow) and genes significantly increased in translation efficiency by 24 hours of SARS-CoV-2 infection in Calu3 cells (green).

dsRNA in virally infected cells can trigger the activation of PKR leading to sustained phosphorylation of EIF2A and inhibition of translation initiation thereby limiting the availability of translation machinery for the virus [23]. A subset of (∼200 genes in mouse embryonic fibroblasts) stress-response genes are resistant to phosphorylated EIF2A-induced shut down owing to the presence of upstream open reading frames (uORFs) in their 5’UTRs [17, 24]. To determine if uORFs conferred protection from translation inhibition seen here, we sorted genes based on the presence of an upstream open reading frame (uORF) in the 5’ UTR as catalogued for 2659 human genes by McGillivary et al. [25]. JUN, ZBTB20 and IL6 all have no uORFs, while 1.5% of downregulated genes contained no uORF and 1.8% containing at least one uORF, respectively were decreased in translation efficiency (Figure 6C). Across all genes the mean log2 fold change in translation efficiency was lower for those genes containing an uORF (one-way analysis of variance [ANOVA], *p* < 0.001; Figure 6D. Since mRNAs predicted to contain at least one uORF are more likely to be decreased instead of increased in translation efficiency, uORFs are protective against translation inhibition by SARS-CoV-2 infection. To further explore this, we obtained a list of 68 homologous genes known to increase in translation during EIF2A phosphorylation (EIF2A-P) in mouse embryonic fibroblasts and contain an uORF in humans. We then compared them with significantly altered genes from our mock versus SARS-CoV-2 infected comparison of translation efficiency. We found no EIF2A-P enhanced genes in the genes upregulated in translation efficiency by SARS-CoV-2 infection (Figure 6E). Eighteen genes were downregulated in translation efficiency by SARS-CoV-2 but known to increase in the presence of EIF2A-P in mouse embryonic fibroblasts. This finding gives further weight to the hypothesis that downregulation of translation efficiency by SARS-CoV-2 is independent of EIF2A-P.

## DISCUSSION

Largely driven by signaling molecules called interferons (IFNs), the innate immune system first recognizes non-self, pathogen-associated molecular patterns (PAMPs) such as double stranded RNA through host proteins DDX58 (RIG-I) and toll-like receptor 3/7 (TLR3/7). This recognition initiates phosphorylation and activation of transcription factors IRF3, AP1 complex and NFKB. In infected lung epithelial cells, these transcription factors activate transcription of genes which can be categorized into three groups. i) Type I and III IFNs (IFNB and IFNL) which increase transcription of interferon stimulated genes (ISGs) including IRF7 which fuels further IFN transcription [26], ii) Proinflammatory cytokines TNF, IL-6 and IL-1β which prepare bystander immune cells to fight the infection and iii) chemokines like CCL5 (RANTES) and CXCL10 (IP-10) to attract distal immune cells to the site of infection. IFNs are arguably the most immediate of these three antiviral effectors, binding to their cognate receptors on already infected and neighboring cells, activating kinases JAK and TYK2, which then phosphorylate STAT1, STAT2, MAPK, PI3K allowing their translocation to the nucleus for activation of target genes with diverse antiviral functions [26].

SARS-CoV-2 is remarkably sensitive to type I IFN pre-treatment [11], suggesting that limiting IFN production, rather than function is the main mechanism through which SARS-CoV-2 achieves high viral loads concomitant with delayed or absent symptoms. Non-structural protein 1 (Nsp1) was found to be one of the most potent inhibitors of IFNB1 promoter induction of 26 viral proteins tested [27]. Nsp1 acts to inhibit host translation by blocking the mRNA entry tunnel on the 40S ribosome [28]. This interaction can also lead to endonucleolytic cleavage in the 5’ UTR preventing the recruitment of more ribosomes to host mRNAs [29, 30].

Studies in bronchial epithelial cells show posttranscriptional restriction of IFNL1/2, IFNB1, CCL5 and IL6 by SARS-CoV-1 and not Dhori virus, an orthomyxovirus known to also productively infect 2B4 cells [31]. Here we corroborate these findings in Calu3 cells using an unbiased genome-wide methodology showing that antiviral cytokines (IFNL1-3, IFNB1, CCL5, CXCL11 and IL6) are restricted in translation at 24 hours post infection. This is consistent with the delayed IFN response seen in COVID19 [32]. Importantly, in a mouse model of SARS-CoV, IFN was only detected in the lung after viral load had already peaked rendering this arm of innate immunity entirely ineffective [33].

A host driven delay in IL6 protein production is unlikely because poly I:C treatment of Calu-3 cells results in a 2-12 fold increase in secreted IL-6 in just 4 hours, with the upper range dependent upon a Th1/17 cytokine environment [34]. IL6 is a significant driver of fever [35], which was an initial symptom in 41% of 174 health care workers that tested positive for SARS-CoV-2 infection upon routine screening [36]. Delayed IL6 production could explain the relatively slow onset of symptoms in SARS-CoV-2 infection. Medium time to symptom onset is 5.1 days and up to 14 days [37], compare to influenza A 1.4 days, influenza B 0.6 days, rhinovirus 1.9 days, parainfluenza virus 2.6 days, non-SARS human coronavirus 3.2 days and respiratory syncytial virus 4.4 days, [38]. Moreover, early IL6 and CCL5 signalling enhances recruitment of innate immune cells and resolution lung pathology caused by respiratory syncytial virus [39, 40] suggesting that timing is important for the antiviral activity of these pro-inflammatory cytokines.

In addition to the abovementioned antiviral cytokines, we show that select transcription factors are restricted at the posttranscriptional level, with JUN, ZBTB20, ATF3, HIVEP2 and EGR1 showing the greatest restriction when accounting for the increase in substrate mRNA while REL and CREB5 were significantly upregulated by RNA-seq but not Ribo-seq when analysis was performed on un-normalized Ribo-seq reads. JUN regulates transcription of IFNB1 mRNA as part of the enhanceosome, encompassing ATF2, JUN, IRF3, p50, p65, CBP and p300 [41]. Post-transcriptional restriction of JUN could limit enhanceosome formation and IFN induction. Indeed, weak induction of IFNB1 mRNA was seen by SARS-CoV-2 in Calu-3 cells compared to Sendai virus [42]. ZBTB20 promotes Toll-like receptor innate immune responses by inhibiting transcription of IKBA [43], decreased protein expression of ZBTB20 may therefore delay immune responses.

In contrast, ATF3 is anti-inflammatory, reducing IL6 and IL12B transcription as part of a negative feedback loop [44]. Restriction in ATF3 protein expression may contribute to un-controlled IL6 production in cells that overcome the translational block, either through competition with ever increasing amounts of IL6 mRNA [45] or through an increase in IL6 mRNA stability enacted by the balance of RNA binding proteins ZC3H12A (Regenase-1) and ARID5A [46]. The latter of which stabilizes IL6 and is negatively regulated by MAPK14 (p38 MAPK) to resolve inflammation [47]. The increase in MAPK14 (p38 MAPK) signalling in aged tissues may contribute to uncontrolled IL6 production [48]. HIVEP2 may also be anti-inflammatory, acting as an NFKB inhibitor, enabling cell survival during memory T cell development [49]. Whether HIVEP2 translational restriction contributes to immune dysregulation or enhances SARS-CoV-2 in infected airway epithelial cells is unclear.

EGR1 does have a clear antiviral function, suppressing foot-and-mouth disease virus (FMDV), Sendai virus and Seneca Valley virus through phosphorylation of TBK1, an important kinase that activates IRF3 [50]. Pos-transcriptional restriction of EGR1 may contribute to delayed IFN responses. Transcription factors REL and CREB5 were significantly upregulated by RNA-seq but not Ribo-seq. REL in heterodimers with RELA is a key transcriptional activator of IFNL1 in human airway epithelial cells [51], thus translational blockage could potentially limit IFNL1 transcription. However, Finally, CREB5 downregulation allows viral persistence in nasopharyngeal epithelia of FMDV-infected animals via a decrease in chemokine transcription suggesting it may also be antiviral [50].

RNA binding proteins, PARP14 and ZC3HAV1 (PARP13) also showed post-transcriptional restriction. Loss of PARP14 strongly attenuates inducible transcription of IFBN1 [52] and Coronaviruses encode a macrodomain protein (Nsp3) that inhibits PARP14 to counteract the host antiviral IFN response [53]. Translational repression of PARP14 may further limit this anti-coronaviral protein to support viral replication. ZC3HAV1 plays important roles in the posttranscriptional regulation of mRNA during the stress response enacting degradation of host and viral mRNAs, translational repression, and microRNA silencing [54]. ZC3HAV1 is known to be antiviral to a suite of viruses including retrovirus, alphavirus, filovirus and hepadna virus by degrading viral RNA. It is also pro-apoptotic by inducing degradation of TNFRSF10D (TRAIL Receptor 4) mRNA, a decoy receptor in the TRAIL-induced apoptosis pathway. Thus, post-transcriptional restriction of ZCSHAV1 would likely favour SARS-CoV-2 replication.

Many cytokines, transcription factors and growth factors are time limited in their effects by RNA binding proteins recruited to AU-rich elements (ARE) in their 3’ untranslated region (UTR). Conditions where ARE-containing mRNAs are stabilized for too long can lead to chronic inflammation [55]. For example, decreased expression of the ARE binding protein AUF-1 in chronic obstructive pulmonary disease (COPD) patient samples leads to stabilization of IL-6, CCL2, CCL1 and CXCL8 mRNA [56]. Many of the genes we found to be inhibited in translation were unstable. JUN, ZBTB20, ATF3 and HIVEP2 have half-lives of 1.3, 2.91, 2.17 and 2.03 hours in contrast to IRF7 which has a half-life of 15 hours. In mouse embryonic stem cells IL6 has a half-life of only 1 hour. This was confirmed in lung epithelial cells where half-life was 40 minutes in the presence of TNF and 80+ minutes in the presence of IL-17 [57].

Assuming a closed loop model of translation facilitated by the eIF4G (5’ cap) to poly(A)-binding protein (3’ tail) interaction [58], stabilized mRNAs are more likely to have ribosomes that re-initiate on the same mRNA, a process coined closed-loop assisted reinitation (CLAR) [59]. Mammalian ribosomes translate at approximately 6 seconds per amino acid [60] and must be spaced ∼100 nucleotides apart [61]. Thus, IL6 would only have time to reinitiate approximately four ribosomes per mRNA, HIVEP2 only one, while IRF7 would survive long enough to house its maximum of 15. Consistent with this, mRNAs bound by multiple ribosomes have higher mRNA stability than those bound by a single ribosome [62]. mRNAs using CLAR are less dependent upon scanning mechanisms performed by EIF4F/EIF4A and initiate more efficiently [63]. Mechanisms of translation inhibition occurring prior to mRNA engagement i.e. virally encoded Nsp1 [45] and host encoded 4E-BP1 [64] should be less effective on closed loop mRNAs. While there currently lacks direct evidence that CLAR ribosomes resist translation inhibition by viral Nsp1 or host 4E-BP1, the increased representation of unstable mRNAs in downregulated genes shown here, combined with a potential for reduced access to the mRNA entry tunnel in CLAR ribosomes suggests this might be the case and warrants further investigation.

We found that mRNAs containing at least one high-confidence uORF were more likely to be decreased in translation efficiency. Given that most genes preferentially translated in the presence of PKR phosphorylated EIF2A contain an uORF we propose that PKR is not directing posttranscriptional restriction in SARS-CoV-2 infected Calu3 cells at 24 hours post infection. Moreover, eighteen genes known to be increased in translation in the presence of phosphorylated EIF2A were significantly decreased in translation efficiency by SARS-CoV-2 suggesting a mechanism independent of EIF2A repression. This leaves Nsp1 or 4E-BP1 as likely responsible for the restriction and further experiments will aim to confirm this.

Research into the human immune response to SARS-CoV-2 infection is still in its infancy. We have limited patient data on the early molecular response because when patients present to the clinic, they have been infected for days already, yet this response is so important for viral control and transmission reduction. Here we have used RNA-seq and Ribo-seq in human bronchial epithelial cells to assess the early response to SARS-CoV-2 infection at the level of transcription and translation. We found a robust antiviral host response at the level of transcription with upregulation of IFNB1 and IFNLs, Interferon-induced proteins with tetratricopeptide repeats, oligoadenylate synthase, antiviral cytokines and transcription factors following SARS-CoV-2 infection of Calu-3 cells, strengthening previous findings [65]. The biological effects of these genes are mediated by protein not mRNA. We have increased the biological relevance of transcriptome analysis by demonstrating restricted translation of IFNB1, IFNL1-3, CCL5, CXCL11, IL6, JUN, ZBTB20, ATF3, HIVEP2, REL, PARP13, PARP14 and EGR1, which have direct antiviral and/or immune regulatory functions. Moreover, we found that unstable mRNAs are more sensitive to SARS-CoV-2 induced translation inhibition and propose that the inhibition may occur at the level of mRNA engagement with the ribosome. Upstream open reading frames were not protective suggesting a mechanism independent of EIF2A repression, which incidentally occurs post mRNA engagement.

In summary, we identify a selection of antiviral and immunological genes that are restricted early in SARS-CoV-2 infection of human cells and highlight that mRNA stability contributes to gene selectivity. Since cytokine mRNA stability is altered in chronic inflammation, assessment of translation restriction during SARS-CoV-2 infection in these contexts is warranted. This may help us understand the wide range of prognoses during infection. Presently, this study provides detailed molecular insight into the delayed interferon response employed by SARS-CoV-2.

## MATERIALS AND METHODS

### Cell culture

VeroE6 cells (ATCC CRL-1586) were maintained in Gibco Dulbecco’s Modified Eagles Medium (DMEM) supplemented with 10 % (v/v) foetal calf serum (FCS), 100 U/mL penicillin, and 100 μg/mL streptomycin (Life Technologies). Calu3 cells were maintained in Gibco Modified Eagles Medium (MEM) supplemented with 20 % (v/v) FCS, 10 mM HEPES, 0.1 mM non-essential amino acids, 2 mM glutamine, 1 mM sodium pyruvate, 100 U/mL penicillin, and 100 μg/mL streptomycin (Life Technologies). All cells were kept at 37 °C in a humidified incubator (5% CO2).

### Infections

All virology work was conducted at the CSIRO Australian Centre for Disease Preparedness at physical containment (PC)-4. The isolate of SARS-CoV-2 (BetaCoV/Australia/VIC01/2020) was received from the Victorian Infectious Disease Reference Laboratory (VIDRL, Melbourne, Australia) and passaged in VeroE6 cells for isolation, followed by passaging in VeroE6 cells for stock generation. All virus stocks were aliquoted and stored at −80°C for inoculations. The infectious titres of SARS-CoV-2 stocks was determined by TCID50 assays performed as described previously [66]. Samples were titrated in quadruplicate in 96-well plates, co-cultured with VeroE6 cells for four days and monitored for development of cytopathic effects (CPE). Calu3 cells were fixed for 30 min in 4 % paraformaldehyde (PFA) and stained with a polyclonal antibody targeting the SARS-CoV-2 Nucleocapsid (N) protein (Sino Biological, catalogue number: 40588-T62, used at 1/2,000) for 1 h. Cells were subsequently stained with 1/1,000 dilution of an anti-rabbit AF488 antibody (Invitrogen catalogue number A11008). Nuclei were counter-stained with diamidino-2-phenylindole (DAPI). Calu3 cells were imaged using the CellInsight quantitative fluorescence microscope (Thermo Fisher Scientific) at a magnification of 10 x, 49 fields/well, capturing the entire well. The relative viral antigen staining was quantified using the Compartmental analysis bioapplication of the Cellomics Scan software. Calu3 cells were grown to 80 % confluency in T25 culture flasks prior to infection with SARS-CoV2 (MOI 1) or mock infected for RNA sequencing experiments.

### MNAse digest and RNA extraction

Isolation of whole-cell RNA and purification of ribosome-protected fragments (RPFs) was performed based on the protocol described in (Reid *et al*, 2015, Methods). Cells were harvested on a dry ice-ethanol slurry in 300 μL ice-cold lysis buffer (1 %(v/v) IGEPAL, 200 mM KOAc, 25 mM K-HEPES pH 7.2, 10 mM MgCl, 4 mM CaCl_2_) and incubated on ice for 10 mins. The lysate was then clarified by centrifugation (8600 xg, 5 mins, 4°C). 50 μL clarified lysate (supernatant) was set aside for RNA-seq analysis while 200 μL was digested with 300 U/mL micrococcal nuclease S7 (MNase; Sigma Aldrich) for 30 mins at 37°C to generate ribosome-protected fragments (RPFs). Both digested and undigested RNA was then purified by phenol:chloroform extraction using TRI reagent® (Sigma Aldrich). The air-dried pellet containing undigested RNA was resuspended in nuclease-free water prior to RNA sequencing.

### Phosphatase treatment and size selection

The air-dried pellet containing the extracted MNase-digested RNA was treated with T4 polynucleotide kinase (Promega) to generate compatible ends for sequencing. RPFs were purified from undigested RNA via electrophoretic separation on a TBE-urea polyacrylamide gel (ThermoFisher) and excision of the region between 25 and 35 nt using an RNA marker. 400 μL 400mM NaOAc, pH 5.2, was added to the excised gel and incubated for 10 mins at −80°C. RPFs were then purified from the gel through three successive cycles of incubation at 95°C for 5 mins followed by vortexing for 20 mins, before extraction using a 0.45 μm cellulose acetate column (Corning) by centrifugation (10 mins, 20,000 xg). 1 mL ethanol and 30 μg/mL GlycoBlue™ co-precipitant was then added to the flowthrough and incubated for 1 hour at −20°C to precipitate the RPFs, followed by centrifugation (15 mins, 20,000 xg, 4°C). The resultant pellet containing RPFs was washed twice in ice-cold 80% (v/v) ethanol, with centrifugation (5 mins, 20,000 xg, 4°C) following each wash. The RPF pellet was then air-dried and resuspended in nuclease-free water prior to RNA sequencing. The quality and quantity of RNA was assessed for all samples using a Bioanalyzer (Agilent, Santa Carla, USA).

### RNA sequencing

RNA-Seq was performed by the Australian Genome Research Facility (AGRF). Illumina TruSeq Stranded mRNA and small RNA libraries were prepared, followed by sequencing on an Illumina Novaseq-6000. Raw data were assessed for overall quality using fastqc v0.11.8. (http://www.bioinformatics.babraham.ac.uk/projects/fastqc/).

### Bioinformatic analysis of RNA-seq reads

Quality and adapter trimming was performed using TrimGalore v0.6.4 (http://www.bioinformatics.babraham.ac.uk/projects/trim_galore/) with default settings for automatic adapter detection. Adaptor trimmed and quality filtered reads were mapped to the human genome (GENCODE v35 primary assembly of GRCh38.p13) using STAR aligner (version 2.5.3a) [67]. Reads mapping to coding sequence (CDS) were counted to quantify transcripts capable of being translated into protein using featureCounts [68]. The Bioconductor package DESeq2 package in R (version 3.6.3) was used to test for differential expression between different experimental groups [69].

### Bioinformatic analysis of Ribo-seq reads

Adaptor was trimmed from the reads using Cutadapt followed by filtering for quality using FastX-toolkit as appropriate for small read analysis. Trimmed and filtered reads were then mapped using Bowtie version 1.2.2 (http://bowtie-bio.sourceforge.net/manual.shtml) to the SARS-CoV-2 genome (accession number MT007544.1), human protein coding transcripts downloaded from BioMart (https://m.ensembl.org/info/data/biomart/index.html), miRNAs from miRbase (http://www.mirbase.org), rRNA from the silva database (https://www.arb-silva.de), snoRNA from the snoRNA-LBME-db database and tRNA from the GtRNAdb database [70]. Percentage of reads with at least one reported alignment were collected from alignment outputs and plotted using ggplot2 in R. Reads mapping to non-coding RNA were filtered out and mapped to the human genome (GENCODE v35 primary assembly of GRCh38.p13) using STAR aligner as per RNA-seq reads with the following parameters --outFilterMismatchNmax 2 --quantMode TranscriptomeSAM GeneCounts --outSAMattributes MD NH –outFilterMultimapNmax 1. Reads mapping to CDS were counted and differential expression analysis run as per RNA-seq. For gene feature mapping statistics, Samtools version 1.10.0 [71] was used to generate mapping stats on reads aligned to the indicated features (again obtained from BioMart) after filtering out reads mapping to non-coding RNA. The exception was stop and start codon sequence which was obtained using the STAR alignment files followed by featureCounts as per CDS counts but using stop_codon for the -t flag. For coverage analysis, transcriptome-mapped bam files from the STAR alignment were sorted and indexed using Samtools. Sorted bam files were converted to bed files and read depth at each genome position with 1-based coordinates determined using Bedtools version 2.29.2 [72] using a library normalisation factor obtained from DESeq2 analysis of the different experimental groups. For metagene analysis of coverage relative to the start codon, the distance from the transcription start site for each gene was extracted from the GENCODE v35 primary assembly of GRCh38.p13 gtf file then used to summarise coverage for across genes using scripts in R. Read-length distributions were obtained using Samtools stats on position sorted alignment files from reads mapped to human CDS and 3’UTR sequences downloaded from Biomart (https://m.ensembl.org), read-length distributions were summarised across all genes. Differential expression analysis of translation efficiency was performed using Riborex package in R using DESeq2 as the modelling engine [21].

## AUTHOR CONTRIBUTIONS

Conceptualization, M.R.A., C.R.S.; methodology, M.R.A., P.J.V.V., A.M.B., C.R., C.R.S., C.C.; software, M.R.A., P.J.V.V., C.C.; validation, M.R.A., A.M.B., C.R., C.R.S. and L.T..; formal analysis, M.R.A., P.J.V.V.; resources, A.G.D.B., C.R.S.; data curation, M.R.A., A.M.B., P.J.V.V.; writing—original draft preparation, M.R.A.; writing—review and editing, P.J.V.V., A.M.B., C.R.S., L.T., A.G.D.B.; visualization, M.R.A., P.J.V.V., C.C, A.M.B., L.T., C.R.S..; supervision, C.R.A, A.G.D.B.; project administration, X.X.; funding acquisition, A.G.D.B. and C.R.S. All authors have read and agreed to the published version of the manuscript.

## FUNDING

This work was funded by the CSIRO. L.T., M.R.A. and A.M.B. are the recipients of CSIRO CERC post-doctoral fellowships.

## DATA AVAILABILITY

All raw data are deposited on the Sequence Read Archive (SRA) under BioProject ID:PRJNA704763. R code used in the analysis of sequencing reads and in making the plots is available here https://github.com/Marina0729/SARSCOV2_Ribo-seq.

## ACKNOWLEDGEMNTS

We are grateful for support from our colleagues at the Australian Centre for Disease Preparedness (https://www.grid.ac/institutes/grid.413322.5) for providing the facility used in the completion of this work. We also thank Shuning Shi, Kerri Bruce and Tamara Gough for maintenance of cell culture. Ondrej Hlinka in CSIRO Information, Management and Technology Business Unit for assistance in Sci-entific Computing.

## Notes

### Competing Interest Statement

The authors have declared no competing interest.

